# A Novel Localized Tracing Technique to Explore Intra-Amygdala Functional and Structural Connectivity Patterns as Mediators of Individual Variability in Stress Response

**DOI:** 10.1101/2022.11.29.517584

**Authors:** Allie Lipshutz, Victoria Saltz, Kristin R Anderson, Alessia Manganaro, Dani Dumitriu

## Abstract

Neuropsychiatric disorders including anxiety and depression can be induced by stress, but not all individuals exposed to stress develop psychopathology. Therefore, probing the neural substrates that underlie trait vulnerability to stress may open the door for preventive approaches that use biological markers to identify at-risk populations. Here, we developed a novel tracing technique to probe local connectivity patterns as predictors of individual variability in stress responses. Specifically, we combined a retrograde transsynaptic rabies tracing system with cFos colocalization immediately after an acute stressor to elucidate local structural and stress-activated (functional) differences in connectivity within the amygdala complex in female and male mice along a spectrum of social approach/avoidance following acute social defeat stress. While we find no structural or functional connections within the amygdala complex as predictors of individual variability in our behavioral readout, our methods provide a novel approach to investigating cellular and behavioral individual variability in stress responses. Furthermore, we identify overall stress-activation as a predictor of social approach/avoidance in two subregions, medial amygdala and piriform-amygdala area, which may serve as potential biological markers of trait vulnerability, with possible clinical applicability.

**Significance Statement:** Exposure to stress is ubiquitous, but the outcomes of stress exposure vary greatly between individuals. Our work introduces a novel tracing technique to probe for structural and functional connectivity patterns across a broad spectrum of behavioral responses to social stress. Utilizing pre-clinical classification in conjunction with representative behavioral classification introduces to the field a mechanism to identify potential clinical targets for preventative screening for neuropsychiatric disorders as well as further individualized treatment. Notably, our work identifies two intra-amygdalar neural correlates of social stress, opening the door for future investigation of the role these regions play in mediating social stress.

## Introduction

Neuropsychiatric disorders, including anxiety and depression, place an enormous burden on the 21% of the population directly affected^1^, as well as on surrounding society. Since 2008, the United States has invested over 6.5 billion dollars of federal funding towards depression research^2^. While progress and discoveries have been made, still over a third of those suffering from depression do not respond to the current available treatment options^3, 4^. Notably, 82% of those with treatment-resistant neuropsychiatric disorders were found to have at least one relevant pre-existing condition^4^. While neuropsychiatric disorders often develop in response to a set of triggering stressors^5, 6^, it is crucial to account for individual variability in psychopathological outcomes following stress. Identifying biological markers of susceptibility to neuropsychiatric disease, thus, may open the door for screening protocols and preventative treatment approaches, prior to the onset of stress.

Due to its critical role in the systemic stress response^7–9^, the amygdala complex may be a promising target for identifying pre-existing structural and functional differences that mediate individual variability in the behavioral stress response. Structurally, the amygdala is composed of 13 subnuclei^10^, intermingled with functionally specific subpopulations of neurons. Each neuronal type has notable extrinsic connectivity with anxiogenic and fear learning-associated regions, including the ventral hippocampus (vHPC)^11–13^, nucleus accumbens (NAc)^10, 14^, prelimbic cortex (PL)^15^, and infralimbic cortex (IL)^16–18^. Beyond extrinsic connections, these nuclei also have critical intrinsic connectivity: sensory input enters through the lateral amygdala (LA) which projects to the central lateral amygdala (CeL) and basolateral amygdala (BLA). The CeL, specifically, contains multiple inhibitory circuits with opposing functions that serve to mediate a range of responses, and with the BLA, project to the centromedial amygdala (CeM), and finally to the brain stem to initiate a fear response^19–21^. While long-range connectivity with the amygdala complex in the context of social stress has received significant attention^15, 22, 23^, the local intra-amygdalar connectivity patterns in this behavioral context remain largely unknown.

Our methods seek to elucidate local connectivity and activity within the amygdala complex that is both preexisting of any stressor (structural) and/or associated with acute stress (functional) to assess whether intra-amygdalar connectivity and activity map onto the individual variability in behavioral responses to social stress. To identify local connectivity and activity, we implemented the well-validated rabies system as a retrograde transsynaptic tracer^24^ and a model of acute social defeat stress (ASDS) that allows for preclinical classification of mice along a spectrum of stress vulnerability (ranging from social avoidance to approach behavior) one hour after exposure to six minutes of social stress^15^, without inducing long-term structural changes to the brain. Immunohistochemical staining for the neural activity dependent protein, cFos^25, 26^, was performed to colocalize stress-activated cells and G-deleted pseudorabies-positive cells to determine projection neurons activated by acute stress. While no differences in the structural and functional connectivity patterns of projection neurons correlated with behavior, we identified regional activation patterns in the medial amygdala (MeA) and piriform-amygdala area (PAA) that correlate with social stress avoidance. Critically, we propose a novel methodological approach to probe differential regional activation as it correlates with behavior and to assess the viability of a regional complex as a marker for susceptibility to stress.

## Methods

### 0.1 Animals

Experimental C57BL/6J female and male mice aged 8 weeks were group housed with five mice of the same sex in each cage. All cages were maintained on a 12-hour light/dark cycle (lights on 7:00 am), with food and water ad libitum. All surgical and behavioral experiments were performed during light hours and began when mice were aged 10 weeks. Aggressive retired breeder CD1 male mice from Charles River Laboratories were used for all social defeat experiments and nonaggressive male CD1 mice were used for Social Interaction (SI) testing. All experiments were conducted in compliance with National Institutes of Health Guidelines for Care and Use of Experimental Animals and approved by the Institutional Animal Care and Use Committees at Columbia University Irving Medical Center and the New York State Psychiatric Institute.

### 0.2 Stereotaxic surgery

Mice were injected with mCherry-tagged TVA and G-protein containing virus AAV8-CMV-TVA-mCherry-2A-0G (0.5-0.9 uL, Salk Institute California) into the BLA bilaterally (from Bregma: AP −1.03, ML ±3.34, DV −4.89, angle 0°) under a combined isoflurane anesthetic (2-5%, SomnoSuite, Kent Scientific) and meloxicam analgesic (3 mg/kg) protocol (**Figure 1A**). Mice were allowed to recover and fully express the virally delivered mCherry-tagged TVA and G-protein complexes for three weeks. GFP-tagged glycoprotein-deleted pseudorabies virus, AAV8-3’-N-P-M-eGFP-L-5’ (0.5-0.9 uL, Salk Institute California), was then injected into the BLA of each hemisphere (from Bregma: AP −1.03 mm, ML ±4.43 mm, DV −5.09 mm, angle 11°) (**Figure 1B**) and mice were given a week prior to behavioral testing to allow for full recovery and expression of the GFP-tagged glycoprotein-deleted pseudorabies virus. The dual angles employed for the two sequential injections was developed to ensure high specificity of injection localization in a deep structure. Post-hoc histological confirmation of BLA targeting was performed on all brains and only those meeting both viral localization and behavioral classification criteria were included in further analysis.

**Figure 1.**
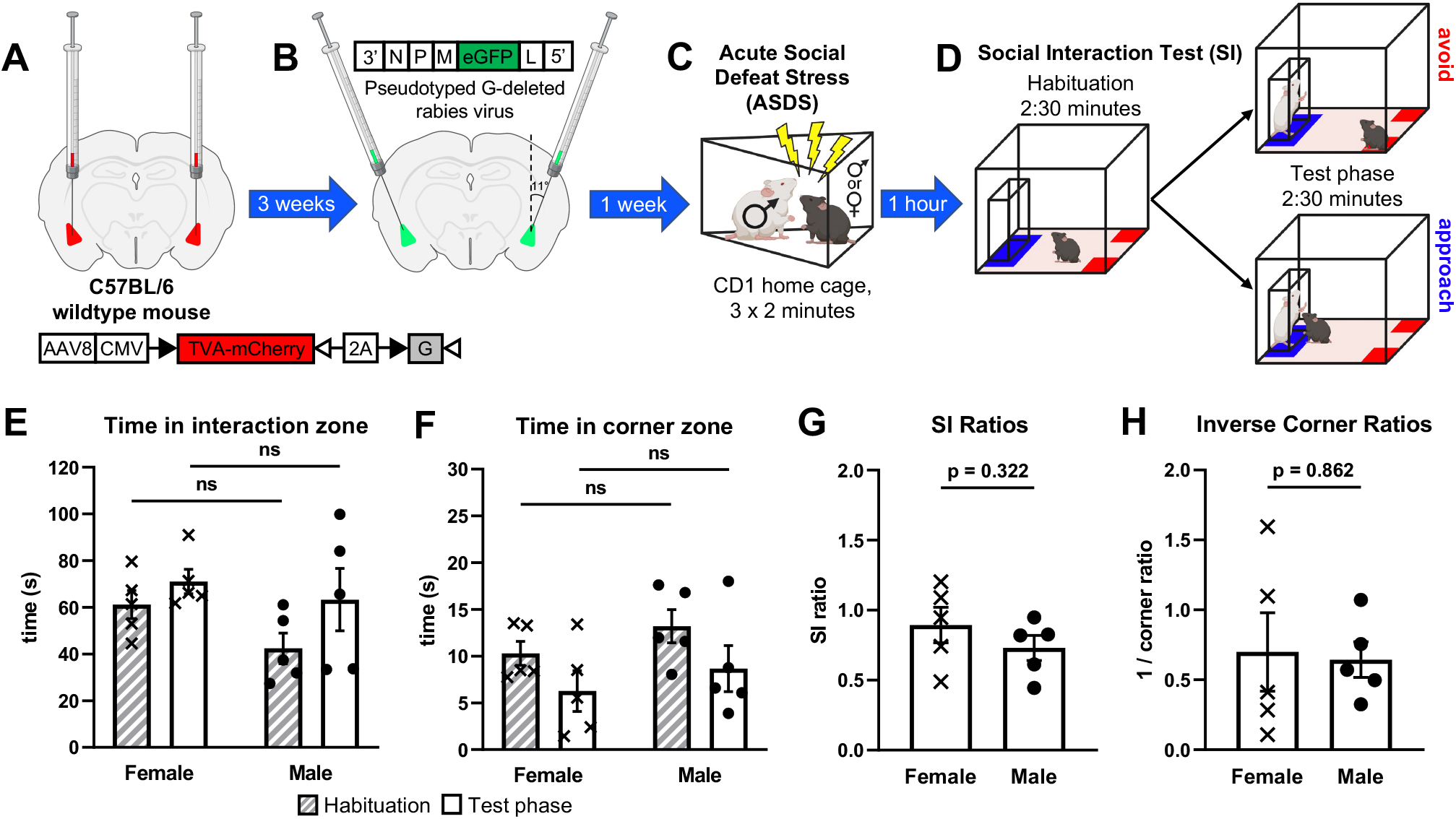
Behavioral responses to acute social defeat stress (ASDS) in female and male mice stereotactically injected for local connectivity tracing in the amygdala complex. **(A)** An mCherry-tagged TVA and G protein-containing viral complex was injected bilaterally into the BLA of female and male C57BL/6 wildtype mice. **(B)** Three weeks later, the same mice were injected with a pseudotyped G-deleted rabies virus into the bilateral BLA at an angle of 11° to promote tracing specificity. **(C)** After a one-week recovery and viral expression period, mice were subjected to ASDS. Each female and male mouse was introduced into the cages of three different territorial aggressive CD1 male mice for a period of two minutes each consecutively. **(D)** A social interaction (SI) test was performed one hour after ASDS, consisting of 2:30 minutes of habituation in which the experimental mouse is allowed to explore the arena and 2:30 minutes of testing in which a nonaggressive CD1 mouse (“target”) was placed in the interaction chamber and the experimental mouse was again allowed to explore. Directly following SI, mice were perfused and brains collected for structural and functional connectivity analyses. **(E)** Female and male mice do not differ statistically in raw time spent in the interaction zone during habituation (F-test, p = 0.872; t-test assuming equal variance, p = 0.0673) or test phase (F-test, p = 0.0972; t-test assuming equal variance, p = 0.604), or in **(F)** raw time spent in the corner zone during habituation (F-test, p = 0.538; t-test assuming equal variance, p = 0.221) or test phase (F-test, p = 0.818; t-test assuming equal variance, p = 0.486)). **(G)** SI ratios were calculated as time spent in the interaction zone when target is present divided by time spent in the interaction zone when target is absent and showed no statistical difference between female and male mice (F-test, p = 0.515; t-test assuming equal variance, p = 0.322). **(H)** Corner ratios were calculated analogously and inverted such that higher scores would indicate more social avoidance analogous to SI ratios and also showed no statistical difference between female and male mice (F-test, p = 0.156; t-test assuming equal variance, p = 0.862).

### 0.3 Acute social defeat stress (ASDS)

One hour before beginning ASDS, urine was collected from nonaggressive CD1 mice and thoroughly applied (10-25 uL) to the urogenital region of all female C57 mice as previously described^27^ to increase the propensity of CD1 aggression upon female mice^28, 29^. Each male C57 was placed into the home cage of a CD1 aggressor for 2 minutes. This was repeated three times sequentially with three different CD1 aggressors, with no rest periods. Experimental female mice were maintained in group housing in the experimental room during the CD1-on-male C57 aggression and were then each individually exposed to acute stress in an identical paradigm as to which the males were exposed (**Figure 1C**). Directly preceding exposure to the first CD1, 20 uL of non-aggressive CD1 urine was applied to the urogenital region of each female C57 mouse. As previously reported^15^, all mice were singly housed for the remainder of the hour and tested for SI 60 minutes following the onset of ASDS. For this experiment, no control C57 mice were included, as the focus of the study was individual variability in stress response.

### 0.4 Social interaction test (SI)

C57BL/6J mice were placed into an open-field arena (45 cm × 45 cm) with a removable wire-mesh enclosure secured in clear Plexiglas (10 cm W × 6 cm D × 30 cm H) placed against the middle of one of the inner walls of the arena. Mice were allowed to habituate to the arena for 2.5 minutes. After 2.5 minutes, a novel non-aggressive CD1 “target” mouse was placed in the wire-mesh enclosure. The C57BL/6J mouse was then allowed to explore for another 2.5 minutes (**Figure 1D**). Open-field arenas and wire-mesh enclosures were wiped with a cleaning solution between tests. All tests were conducted during the lights on period (between 13:00 and 19:00) under infrared light conditions. A video-tracking system (Ethovision XT 15, Noldus Information Technology) was used to record the movements of the C57 mouse during the “target absent” and “target present” periods. The total time spent in the “interaction zone,” defined as a 24 cm × 8 cm corridor surrounding the wire-mesh enclosure, and corner zones, 9 cm × 9 cm in the two corners opposite from the interaction zone, was recorded. The SI ratio was calculated by the absolute time spent in the interaction zone (blue, **Figure 1D**) with the target present divided by the absolute time spent in the interaction zone with the target absent. The corner ratio was calculated analogously, as the absolute time spent in the corner zone (red, **Figure 1D**) with the target present divided by the absolute time spent in the corner zone with the target absent^15^, and was then inverted for enhanced comparability with SI scores (i.e. lower scores predict social avoidance and higher scores predict social approach).

### 0.5 Tissue collection and processing

Mice were anesthetized with isoflurane, checked by toe pinch, and sacrificed by transcardial perfusion with 4% paraformaldehyde immediately after SI testing. Perfusions were performed at a flow rate of 9 mL/min for 10 minutes, followed by decapitation and submandibular cuts for increased fixation. Brains were post-fixed for 72 hours, deskulled, and sectioned at 100 μm using a vibratome (speed 8.5, frequency 9; Leica Biosystems), at which point the left hemisphere was marked with a “nick” for hemisphere-specific analysis of injection accuracy. Sectioned brains were stored in 0.9M PBS with 0.5% sodium azide until immunohistochemical analysis was performed. Sections were serially stained with rabbit anti-cFos (9F6) primary monoclonal antibody (Cell Signaling, catalog number: 2250S, lot number: 12, 1:2000 in 0.3% PBST + 5% Normal Goat Serum + 1% BSA + 0.2% Cold Water Fish Skin Gelatin block solution) and Alexa Fluor-conjugated AffiniPure goat anti-rabbit secondary antibody (Alexa Fluor-647, Sigma, code number: 111-605-033, lot number: 157045, 1:500 in 0.3% PBS-T). All sections were incubated in 4’,6-diamidino-2-phenylindole (DAPI) for histological identification

### 0.6 Imaging and analysis

A Nikon Eclipse Ti2-E Motorized inverted microscope and the Element Acquisition Software were used for all imaging. Triple or quadruple channel z-stacks were acquired to image GFP, mCherry, CY5, and sometimes DAPI signal and the EDF function of the Element Software applied to collapse the z-stack. For histological confirmation of injection sites, low resolution (4X) 16-bit images of sections spanning the entire anterior-posterior extent of the amygdala were inspected. For projection analysis of cFos, GFP-tagged Rabies, and mCherry-tagged TVA-G cells, higher resolution (10X, and 40X) 16-bit images were acquired at the sites of interest (**Figure 2A**). Further analysis was done on two selected sections for each animal, one corresponding to the anterior part of the amygdala complex (AP −1.2 to −1.8 from bregma) and one to the more posterior part (AP −1.5 to −2.1 from bregma). The nd2 files acquired were imported in Fiji^30^ and modified for counting and reconstruction analysis. Each split channel was smoothed and a seven pixels bg subtraction applied. Before proceeding to counting, non-rigid free-form registration of the selected 100 μm sections was performed on the GFP channel with the open-source WholeBrain Software^31^ in RStudio. Briefly, each image was first matched to a corresponding plate (42, AP −1.34 from Bregma) of the Mouse Allen Brain Reference Atlas CCF v3^32^. The contour of the brain section was segmented out using the autofluorescence of the down-sampled section itself, and a set of correspondence points that align the selected reference atlas plate with the tissue-section was automatically generated by the “thin-plate splines algorithm”. Landmarks such as ventricles were used to manually adjust, add or remove correspondent points when necessary. The two sets of reference points (atlas and tissue-section reference points) constitute a 2D final mapping of each brain region along the antero-posterior axis. The extracted warped contours of the amygdala complex were exported from RStudio and overlapped to the original 16-bit images in ImageJ^33^ with a custom-made macro^34^. GFP Rabies^+^ cells were counted after thresholding the resulting images («analyze particle» plugin). CY5 cFos^+^ cells were counted after applying a white top hat to the images (MorphoLibJ plugin). For colocalization analysis of starter cells (GFP Rabies^+^ mCherry TVA-G^+^) and stress-activated projection cells (GFP Rabies^+^ CY5 cFos^+^), the image calculator was implemented using the AND function. Method validation was performed with the coloc function. The schematic of projections and starter cells for the left and right hemispheres (**Figure 2B-C**) were reconstructed in ImageJ^33^ by overlapping and realigning in a stack the segmented images for the channel of interest over the representative plate from the Paxinos Mouse Brain Atlas 4^th^ edition^35^ in Adobe Illustrator V for final color modifications.

**Figure 2.**
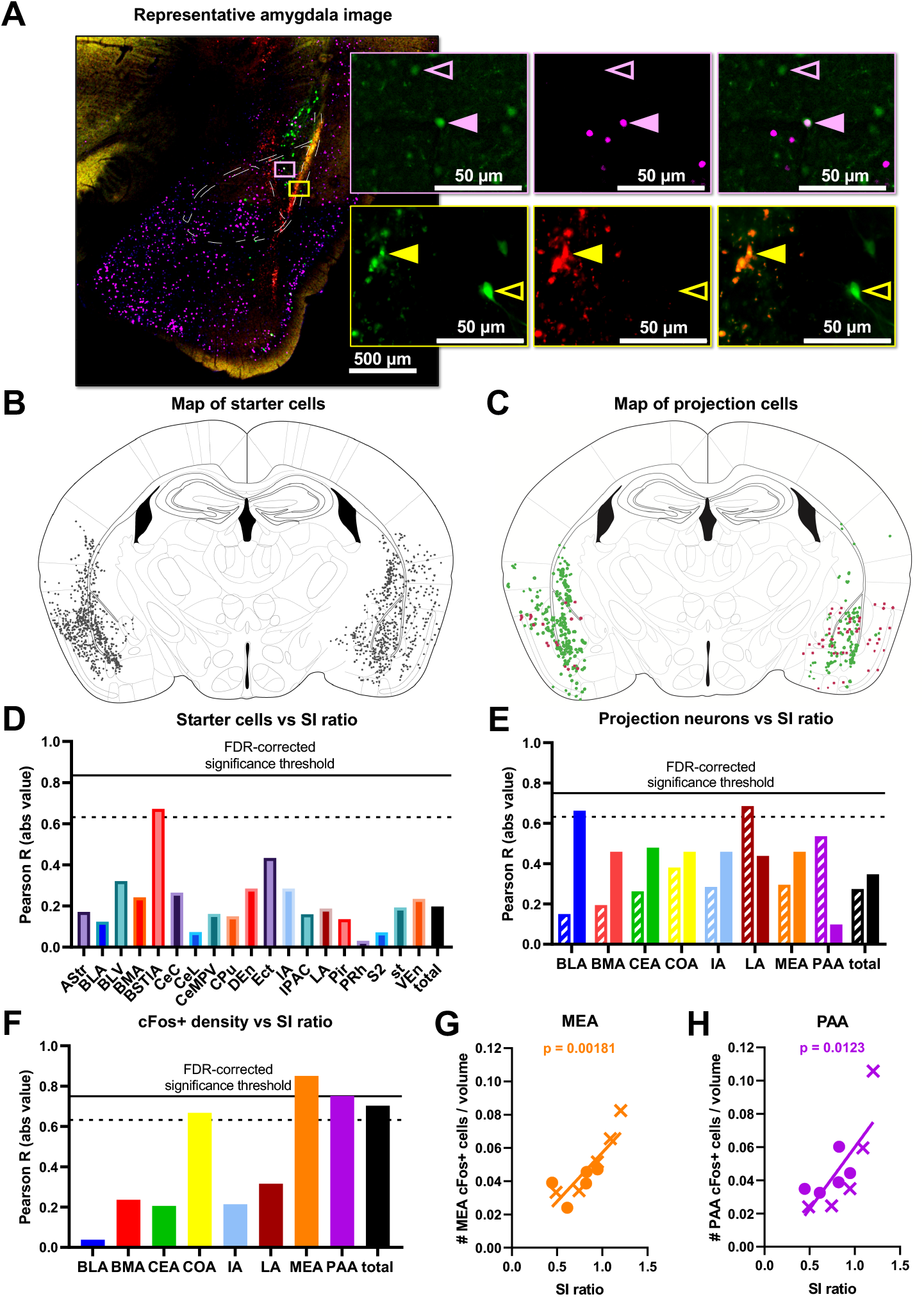
Structural and functional connectivity patterns within the amygdala complex do not mediate individual variability in stress response but MeA and PAA overall activation predicts social approach behavior. **(A)** Example triple-channel image of the amygdala region (red=mCherry, green=GFP, magenta=cFos). Top inset (pink box and outline) shows examples of a structural projection neuron (open pink arrow, GFP^+^/cFos^-^) and a functional projection neuron (closed pink arrow, GFP^+^/cFos^+^). Bottom inset (yellow box and outline) shows examples of a starter neuron (closed yellow arrow, GFP^+^/mCherry^+^) and a projection neuron (open yellow arrow (GFP^+^/mCherry^-^). **(B)** Map of all identified starter cells (GFP^+^/mCherry^+^). **(C)** Map of all identified structural (GFP^+^/cFos^-^) and functional (stress-activated) projection (GFP^+^/cFos^+^) neurons. **(D)** Correlations between starter cell count in each region and SI ratio with FDR-corrected and non-FDR-corrected significance threshold indicated (solid horizontal line, Pearson R = 0.835, p = 0.00263; dashed horizontal line, Pearson R = 0.632, p = 0.05; respectively). Starter cell counts did not predict behavioral phenotype in any identified brain region. **(E)** Correlations between structural projection neuron (GFP^+^/cFos^-^) count (striped) and functional projection neuron (GFP^+^/cFos^+^) count (solid) in each region and SI ratio with FDR-corrected and non-FDR-corrected significance threshold indicated (solid horizontal line, Pearson R = 0.749, p = 0.0125; dashed horizontal line, Pearson R = 0.632, p = 0.05; respectively). No significant correlations identified. **(F)** Correlations between overall number of activated cells per brain region and SI ratios indicate **(G)** MeA and **(H)** PAA activation positively predict social approach (Pearson R = 0.851, p = 0.00181 and Pearson R = 0.751, p = 0.0123, respectively).

## Results

### Behavioral Results

Following adequate recovery time from stereotactic viral injections (**Figure 1A-B**), mice were exposed to our previously developed ASDS paradigm^15^, which consists of three consecutive two-minute bouts of social stress (**Figure 1C**), and was here extended to include female mice. After one hour of rest, the experimental mice underwent an SI test as a behavioral readout of individual variability in behavioral response to stress, measured as time spent in the interaction zone, near a novel “target” mouse of the same breed as the aggressor, and the time spent in the corner zones (**Figure 1D**). Given the explicit readout of individual variability in behavioral response to stress, all mice underwent the social stress paradigm with no stress-controls being included. ASDS elicited a spectrum of behavioral responses ranging from social avoidance to social approach in both female and male mice (**Figure 1E-H**). When compared by a two-sample t-test assuming equal variances female and male mice showed no statistically significant difference on either SI ratios (F-test, p =0.515; t-test, p = 0.322) (**Figure 1G**) or inverse corner ratios (F-test, p = 0.156; t-test, p = 0.862) (**Figure 1H**), providing a justification for concatenating female and male data for a total sample size of 10 animals that could be used for analyses exploring neural correlates of behavioral stress responses.

### Cellular Results

Post-hoc histological confirmation of the injection site was performed on all brains and overlaid with DAPI for regional identification of the amygdala. All ten animals demonstrated injection endpoints of both the mCherry-tagged TVA and G-protein containing viral complex and the GFP-tagged pseudotyped G-deleted rabies virus (**Figure 2B**) within the bilateral amygdala complex. After thorough optimization of immunohistochemical conditions, consistent cFos fluorescence was observed in all ten animal samples. Two amygdala sections from each brain were imaged in detail, one covering the more anterior portion of the amygdala complex (AP −1.2 to −1.8 from bregma) and one covering the more posterior end of the amygdala complex (AP −1.5 to −2.1 from bregma). Starter cells were identified across the amygdala complex as well as surrounding cortical regions in a total of 19 subregions, and starter cell location and total number was not predictive of the behavioral phenotype (**Figure 2D**). Eight of the 13 amygdala subregions contained projection cells (GFP^+^/mCherry^-^), so further analyses were restricted to these eight subregions. By controlling for the individual variability in the infection rate using total starter cell (GFP^+^/mCherry^+^) number, structural (stress-inactive, GFP^+^/cFos^-^) and functional (stress-activate, GFP^+^cFos^+^) projection cells counts were compared across eight amygdala-associated subregions (**Figure 2C,E**). No correlations between structural or functional projection cell number and SI scores in any of the eight amygdala subregions reached FDR-corrected significance threshold. We further explored if the overall activation of these subregions, denoted by cFos^+^ cell density per region (irrespective of GFP^+^ neurons), correlates with SI scores and found the activation in two subregions to reach FDR-corrected threshold of significance: MeA and PAA (**Figure 2F-H**).

## Discussion

Here, we present a novel method for virally tracing localized monosynaptic neuroanatomical connectivity, utilizing the retrograde transsynaptic rabies system and a dual angled injection approach to obtain a higher degree of specificity within a deep structure of interest. While long-range monosynaptic connections to the amygdala are well-described^22, 23^, our methods only resulted in short-range projection tracing. Notably the helper virus used in our experiment, AAV8-CMV-TVA-mCherry-2A-G, has only been used to trace short-range projections to the best of our knowledge^36^. Thus, the combination of utilizing a short-range helper virus serotype, low viral titers, and dual angled injections allowed us to illuminate local projections that had thus far been hidden by viral overload using other viral combinations. Furthermore, for enhanced translatability, our experimental design focuses on a spectrum of SI ratios that more effectively approximates the range of behavioral phenotypes seen in the human clinical setting^37, 38^ than the commonly employed dichotomous classification of “social avoidance” versus “social approach”. Notably, we did not identify any significant correlations between the structural or functional connectivity of projection neurons within the amygdala complex, opening the door to future investigation into other local microcircuits associated with the social stress response.

Though no structural and functional connectivity differences were identified in projection neurons, we do show a significant correlation between overall activation in both the MeA and PAA with SI ratio, suggesting that MeA and PAA activation may be markers of vulnerability to social stress. Structural plasticity in the MeA has been shown to be affected by chronic restraint stress (CRS) in mice, as exemplified by decreased spine density in MeA stellate neurons. Loss of spine density in MeA stellate neurons was further shown to be associated with trait vulnerability to stress and a lower SI ratio, rescuable by acetyl-L-carnitine^39^.

Much like the MeA, the PAA has been largely implicated in the associative learning of sensory cues with valence. Specifically, the PAA has been shown to be active during odor fear learning. Upon optogenetic inhibition of the BLA to PAA circuit, odor fear learning is impaired, suggesting a mediation of odor fear learning by BLA to PAA circuitry. In response to acute stress, mice in our study likely rely primarily on olfactory cues to assess whether to interact with the novel mouse in the social interaction test. By demonstrating that PAA activation is directly correlated with social approach behavior, we suggest that protection against maladaptive behavioral response to stress is associated with, and perhaps reliant upon, effective activation of the odor fear learning network. Inhibition and overexpression experiments of the MeA and PAA will be required to establish a causative link.

The identification of preclinical neural targets in the MeA and PAA, supported by literature, utilize a new classification approach for the social defeat model that more closely mirrors clinical applications. The methodology used here serves as a powerful pilot towards elucidating how connectivity and activity within a localized brain region can be indicative of trait vulnerability to stress. Through the approach of preclinical classification of a social stress response, associating behavioral phenotypes with neuronal correlates, and ultimately identifying target brain regions and circuits that mediate the stress response, we begin to identify potential clinical targets for preventative screening for neuropsychiatric disorders as well as further individualized treatment.

